# Age-Related Balance Problems in Mice Are Sharpened by the Loss of Calcitonin Gene-Related Peptide (CGRP) and a Vestibular Challenge

**DOI:** 10.1101/2023.06.28.546965

**Authors:** Shafaqat M. Rahman, Catherine Hauser, Anne E. Luebke

**Affiliations:** Department of Biomedical Engineering, University of Rochester, Rochester, New York, United States of America; Department of Neuroscience, Del Monte Institute of Neuroscience, University of Rochester Medical Center, Rochester, New York, United States of America

## Abstract

Aging impacts the vestibular system and contributes to imbalance. In fact, in the elderly balance deficits often precede changes in cognition. However, imbalance research is limited in assessing aging mouse models that are deficient in neuromodulators like Calcitonin Gene-Related Peptide (CGRP). We studied the loss of CGRP and its effects in the aging mouse, namely its effect on both static and dynamic imbalances. In addition, postural sway and rotarod testing were performed before and after a vestibular challenge (VC) in the 129S wildtype and the αCGRP (−/−) null mice. Four age groups were tested that correspond to young adulthood, late adulthood, middle age, and senescence in humans. Our results suggest wildtype mice experience a decline in rotarod ability with increased age, while the αCGRP (−/−) null mice perform poorly on rotarod early in life and do not improve. Our postural sway study suggests that a vestibular challenge can lead to significantly reduced CoP ellipse areas (freezing behaviors) in older mice, and this change occurs earlier in the αCGRP (−/−) null mouse. These results indicate that αCGRP is an important component of static and dynamic balance; and that the loss of αCGRP can contribute to balance complications that may compound with aging.

## Introduction

Aging impacts the vestibular system, and contributes to balance problems and an increased risk of falls; for a recent review see [1]. It is estimated that 35% of adults aged 40 and over will experience some form of vestibular impairment, and 85% of people aged 60 and over will experience indications of balance dysfunction [2]. Balance dysfunction causes a nearly 3-fold increase in the odds of falling, and this clinical outcome is highly prevalent in the elderly, affecting 1 in 3 adults aged over 65 per year [3, 4]. In fact, Leach et al. found a linear relationship between day-to-day variability in postural sway and cognitive status in older adults [5], highlighting the deficits that can occur with aging [6–10].

Current knowledge of aging’s effects on human gait and postural control provides context for preclinical researchers to study vestibular dysfunction and aging in mouse models. Mice with loss of calcitonin gene-related peptide (αCGRP) show notable changes in vestibular sensory processing and behavior. The αCGRP (−/−) null mice exhibit a 50% gain reduction in the vestibulo-ocular reflex, shorter vestibular sensory-evoked potentials, and deficits in rotarod ability [11, 12]. Yet, it is unclear if αCGRP loss compounds with aging effects during static and dynamic balance conditions. We hypothesized that αCGRP (−/−) null mice would exhibit severe imbalance phenotypes as a function of aging compared to WT and that a vestibular challenge would aggravate these phenotypes.

We assessed mouse postural sway and rotarod ability on a modified dowel as surrogate behaviors for static and dynamic imbalance in the αCGRP (−/−) null mice and their wild type complement. Mice were studied across four age groups: young adulthood (2.3 to 6 months), late adulthood (6 to 10 months), middle age (10 to 15 months) and near senescence (15 to 18 months). These age groups correspond to 20 to 30 years, 30 to 40 years, 40-50 years, and 50-70 years in humans [13]. In this study, two different vestibular challenges (orbital or circular rotation) were used to perturb mouse behavior during static and dynamic conditions.

## Materials and methods

### Animals

The αCGRP (−/−) null and WT transgenic mice were produced and characterized on a pure 129SvEv background obtained from Emeson laboratory [14]. Mice were shipped to the Luebke laboratory as heterozygous (+/−) and genotyped using previously established protocols. The αCGRP heterozygous (+/−) were bred to generate homozygous αCGRP (−/−) null and WT (+/+) offspring used in these studies. The αCGRP (−/−) null strain has a targeted deletion of αCGRP due to tissue-specific alternative splicing of the calcitonin/αCGRP gene [14]. Mice were housed under a 12 to 12 day/night cycle at the University of Rochester’s Vivarium under the care of the University of Rochester’s Veterinary Services personnel. All procedures have been approved by University of Rochester’s University Committee on Animal Resources (UCAR). A total of 231 mice - 106 wild type (45M/61F) and 125 αCGRP null (59M/66F) - were used for these studies. A table is provided to indicate the number of mice used across the factors of sex, age, and strain, with many but not all mice repeatedly tested as they aged (**Table 1**). Mice were equilibrated in the test room controlled for an ambient temperature between 22-23 ° C for at least 30 minutes prior to testing and remained in this room until the experiment was completed.

**Table 1:**
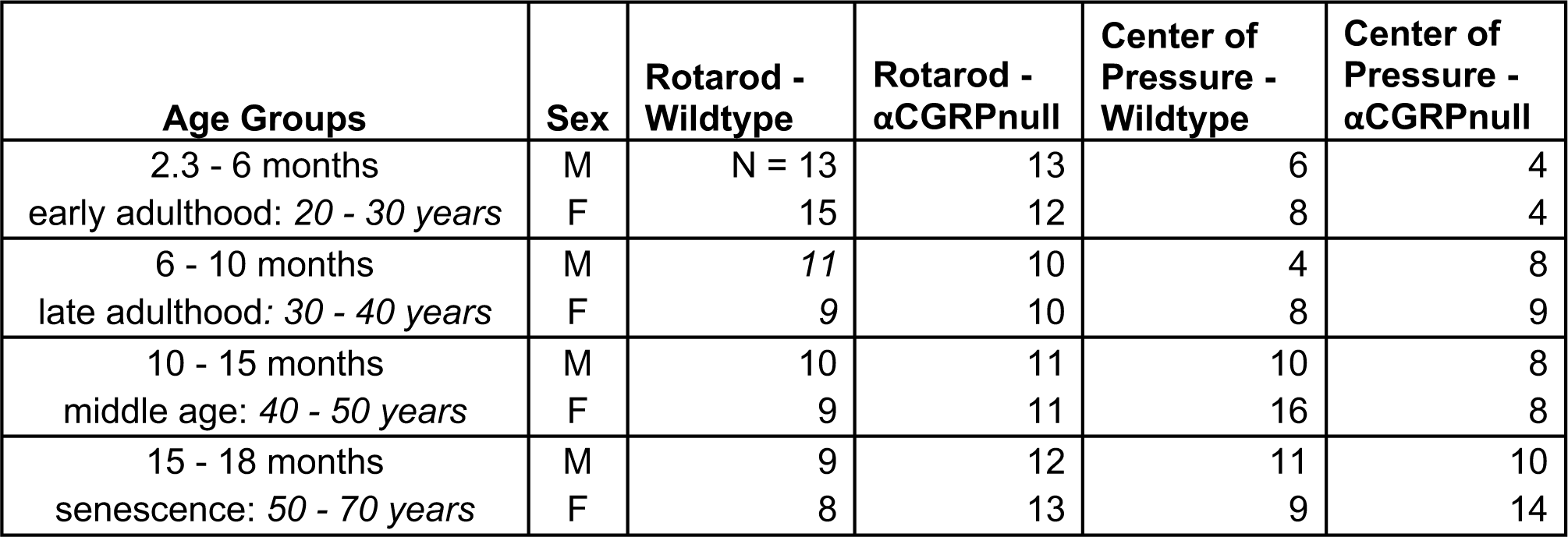
Sample sizes of 129SvEv WT and αCGRP (−/−) null mice are depicted per age group for rotarod and center of pressure tests. Mice were tested in the range of 2.3 – 18 months (many were repeatedly tested to conserve overall animal count) and were categorized into four different age groups: early adulthood, late adulthood, middle age, and senescence. The equivalent age in humans is italicized.

| Age Groups | Sex | Rotarod -<br>Wildtype | Rotarod -<br>αCGRPnull | Center of<br>Pressure -<br>Wildtype | Center of<br>Pressure -<br>αCGRPnull |
| --- | --- | --- | --- | --- | --- |
| 2.3 - 6 months | M | N = 13 | 13 | 6 | 4 |
| early adulthood: 20 - 30 years | F | 15 | 12 | 8 | 4 |
| 6 - 10 months | M | 11 | 10 | 4 | 8 |
| late adulthood: 30 - 40 years | F | 9 | 10 | 8 | 9 |
| 10 - 15 months | M | 10 | 11 | 10 | 8 |
| middle age: 40 - 50 years | F | 9 | 11 | 16 | 8 |
| 15 - 18 months | M | 9 | 12 | 11 | 10 |
| senescence: 50 - 70 years | F | 8 | 13 | 9 | 14 |

### Vestibular Challenge

#### Orbital Rotation

During postural sway testing, mice experienced a vestibular challenge (VC) in the form of a steady, orbital rotation at 125 rpm for five minutes (orbital radius = 2 cm). The mice were placed in an acrylic box attached to an orbital shaker’s surface during rotation, and the mouse’s head is fixed in one direction and does not rotate with the axis.

#### Circular Rotation

During rotarod testing, mice were challenged with circular rotation as the VC. Mice were rotated at 125 rpm at a distance of 7.0 cm from the rotational axis. The mouse’s head is fixed perpendicular to the axis of rotation. This VC is more severe than orbital rotation due to the increased orbital radius requiring the device to transverse a larger circumference to maintain the same rpm, and mice commonly exhibit a transient (1-2 second) body tremor after experiencing the circular rotation that is not seen with the orbital rotation. This VC is only performed for 30 seconds.

### Rotarod testing for dynamic balance assessment

Dynamic balance was assessed with a rotarod (Columbus Instruments) conFig ured with a rat dowel (radius = 3.5 cm). The larger-sized rat dowel was incorporated to facilitate walking and minimize gripping and trapezing behaviors. The rotarod is designed to test four mice simultaneously, and mice were tasked to maintain balance on the rat dowel rotating from 5 to 44 rpm at an acceleration step of 2.4 rpm every 4 seconds. Latency to fall (LTF) is measured when mice fall from the dowel. Two days of rotarod testing were performed. The first day was a training day where mice were tested for 6 to 8 trials. On the second day, mice were briefly trained for 4 trials and were then given a 15-minute rest. After the rest, mice were tested for 3 trials (pre-VC) and were then stimulated with circular rotation as the vestibular challenge (see <u>*Vestibular Challenge – Circular Rotation*</u> for methods). Mice were then immediately tested for 3 trials on the rotarod after the challenge (post-VC). Approximately 10-30 seconds pass in between subsequent trials during pre-VC and post-VC tests. A schematic of the rotarod methods is shown in **Fig 1A**.

**Fig 1:**
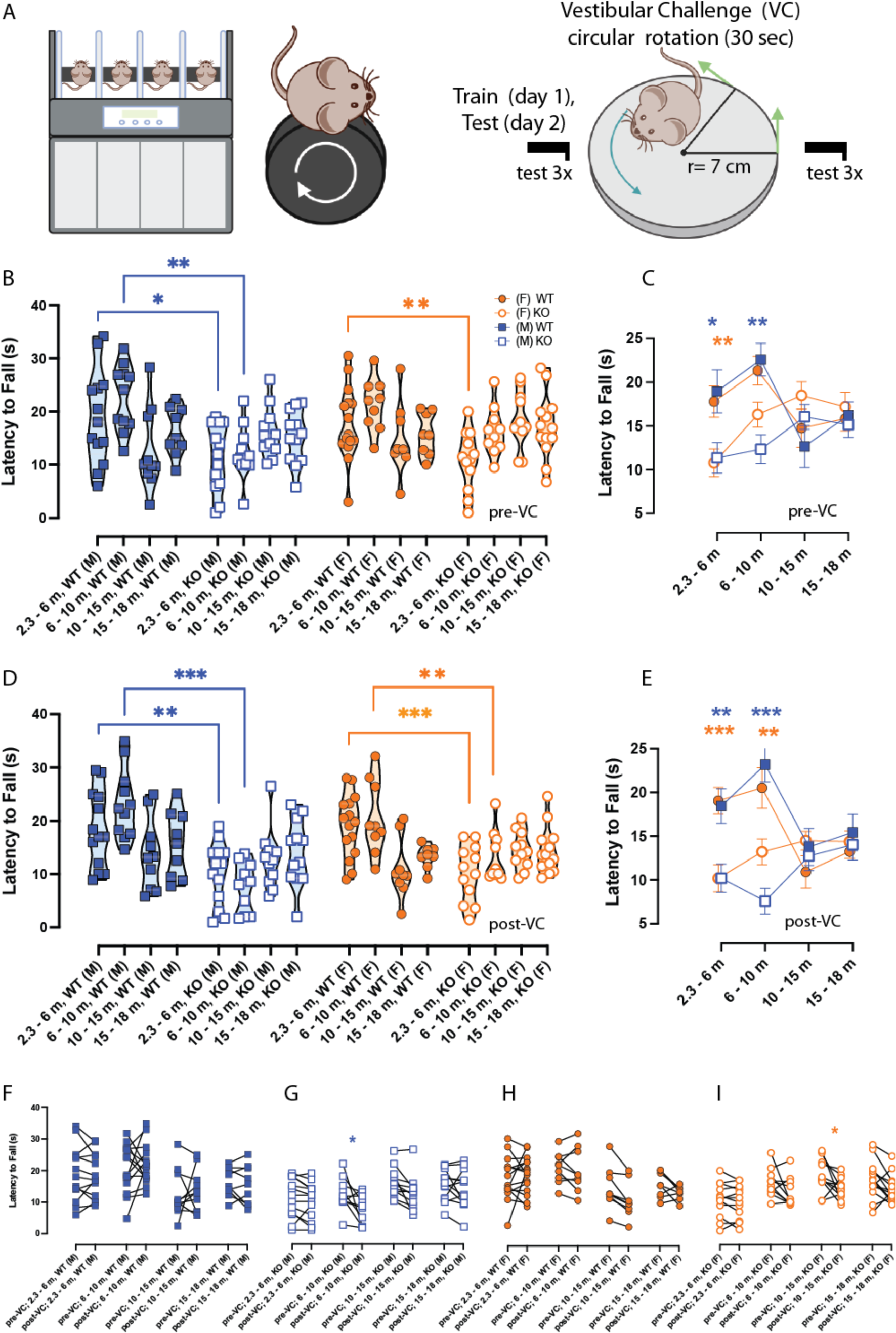
Differences in rotarod performance due to aging, vestibular challenge (VC), and αCGRP loss (−/−). **A**. Mice were tested on rotarod configured with rat dowel (r = 3.5 cm) and were assessed for 3 trials before (pre-VC) and after (post-VC) a 30-second circular rotation (r = 7 cm) across four age groups. Symbols are depicted as follows: female (F) – orange circle, male (M) – blue square, WT – closed, αCGRP (−/−) null (KO) – open. **C, E**. Individual MAX LTFs were grouped and depicted as mean + SEM per age group. **B**. During pre-VC rotarod, WT(M) outperformed KO(M) by 7.6 ± 2.4 s in early adulthood (*p = 0*.*01)* and by 10.3 ± 2.6 s (*p = 0*.*001*) in late adulthood. WT(F) outperformed KO(F) null in early adulthood by 7.0 ± 2.3 s (*p = 0*.*0009*) and appeared to outperform in late adulthood by 5.03 ± 2.8 s, but this was not a significant. WT performs similarly to KO on rotarod after 10 months. **E**. Similar differences with age were seen during post-VC rotarod. In either sex, WT outperforms KO in early and late adulthood but no differences were observed after 10 months. **F-I**. Before-after plots were constructed that highlight the effects of the VC on rotarod, but VC’s effects were generally unclear. Full list of ANOVAs and F-values can be found in top halves of **Table 2 and 3**.

### Postural Sway testing for static balance assessment

The CoP assay is used to measure mouse postural sway as a surrogate for static balance changes [15]. The mice were weighed and placed on a force plate designed to measure the forces due to postural changes in X, Y, Z, and its angular moments in the XY, YZ, and XZ directions (**Fig 2A**). We used the AMTI Biomechanics Force platform (model HEX6×6) and its corresponding AMTI automated acquisition software. An accessory plexiglass cover is placed over the force plate to prevent mice from moving off the force plate. When placed on the force plate, mice are given 2 to 5 minutes to acclimate to the novel environment and minimize their exploratory behavior. After acclimation, 10 trials of the mouse’s CoP area during the pre-VC test were measured by the relative output of four vertical sensors (resolution per CoP trial = 300 samples per second). These trials were captured when the mouse showed no active exploratory behavior (e.g., grooming, standing) and its four paws were touching the surface of the force plate. The mice were then subjected to VC (orbital rotation for 5 minutes) and were then placed back onto the platform. Five minutes after the VC, an additional 10 measurements were measured. A modified MATLAB code was used to analyze CoP data, generating a 95% confidence ellipse that enclosed 95% of the CoP trajectory values computed in a single trial.

**Fig 2:**
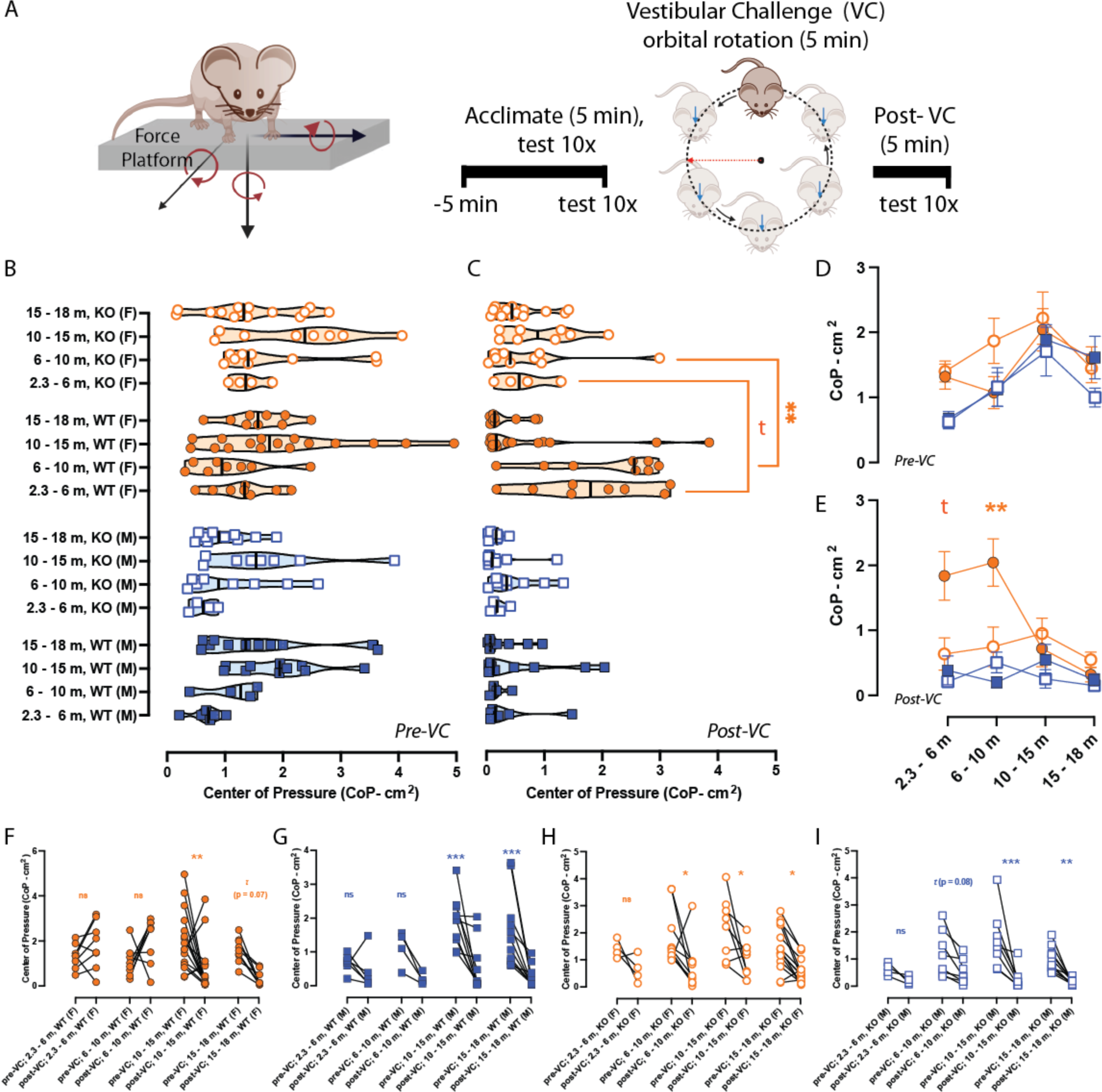
Postural sway (center of pressure – CoP) changes due to aging, vestibular challenge (VC), and αCGRP loss (−/−). **A**. AMTI Biomechanics Force platform (model HEX6×6) and its corresponding AMTI automated acquisition software were used to measure 95% confidence CoP ellipses (cm^2^), and the experimental timeline involves an acclimation period (5 minutes) followed by the experiment where 10 trials are captured before and after a 5-minute orbital rotation as the VC. Symbols and color scheme are depicted like in **Fig 1** previously. **D-E**. Individual average CoPs were grouped and depicted as mean + SEM per age group. **D**. During pre-VC, we observed a general increase in CoP area when examining CoP progressing from early adulthood to middle age, but differences were not seen between strains. **E**. For post-VC, WT (F) had larger CoPs compared to KO(F) in early adulthood (*2*.*3 – 6 m: t (p = 0*.*09)*) and in late adulthood (*6 – 10 m: p = 0*.*009)*, however, this difference is gone after 10 months. **F-G**. Middle-aged and senescent WT(F) and WT(M) exhibit reduced CoP ellipses due to the VC. **H-I**. KO(F) and KO(M) exhibit reduced CoP ellipses due to the VC from late adulthood to senescence, highlighting a VC sensitivity in KO that arises earlier in age than their WT complement. See bottom halves of **Table 2 and 3** for F-statistics and p-values.

### Statistics

Analyses were conducted in GraphPad Prism 9.5. A 10% ROUT removal was used to remove outliers in a given mouse’s test prior to calculating its average CoP. Despite this robust outlier removal, at least six different 95% ellipse areas were used to compute the average. Then, the CoP average (cm^2^) per mouse were further averaged as a mean group CoP + ± SE. For rotarod, data was analyzed by first determining the MAX latency to fall (LTF) from the three trials assessed per testing condition. The MAX LTF values (s) were grouped and computed for an average maximum LTF ± SE. Repeated measure ANOVA (RM-ANOVA) and Bonferroni post-hoc analyses were the primary statistical tool. Significance was established at p < 0.05 for all analyzes.

## Results

### Aging and αCGRP loss (−/−) on rotarod

Prior to the vestibular challenge (pre-VC), mice were trained and later assessed for baseline rotarod ability. Two-way RM-ANOVAs assessed the factors αCGRP loss and aging, and corresponding F-statistics and p-values can be found in the top half of **Table 2**. Unlike WT, the αCGRP (−/−) null mice tested at early adulthood had poor rotarod ability. In **Fig 1C**, we simplified the violin plot in **Fig 1B** by averaging the MAX LTFs per age for each sex and strain. Bonferroni post hoc analyses indicates that WT males outperformed αCGRP (−/−) null males by 7.6 + 2.4 s in early adulthood (*p* = 0.01) and by 10.3 + 2.6 s (*p* = 0.001) in late adulthood. Similarly, WT females outperformed αCGRP (−/−) null in early adulthood by 7.0 + 2.3 s (*p* = 0.0009) and in late adulthood by 5.03 + 2.8 s (though not significant). While WT mice outperform the αCGRP (−/−) null during the first 10 months of age, their rotarod ability declines after 10 months and their performance resemble their similarly aged αCGRP (−/−) null peers. **Fig 1C** suggests that aging can cause poor gait and dynamic balance performance in otherwise healthy controls (WT). However, a mouse lacking αCGRP exhibits poor dynamic balance early on in development and this balance ability does change significantly as they age.

**Table 2:** Separately done in males and female mice, rotarod and postural sway data during pre-vestibular challenge (VC) and post-VC tests were analyzed with two-way repeated measures ANOVA to assess the factors *aging* and *αCGRP loss*. Bonferroni post hoc analyses computed the difference between wildtype and αCGRP (−/−) null at each age group. F-values are listed with respect to degrees of freedom (DF_n_, DF_d_) and p-values are listed accordingly.

| Assay | Biological Sex | ANOVAs<br>F-statistics, p-val | Post-hoc<br>WT vs KO \| p-val |
| --- | --- | --- | --- |
| Rotarod | Male | *Pre-VC*<br>αCGRP loss: $F(1, 81) = 8.30, p = 0.005$<br>Aging: $F(3, 81) = 0.923, p = 0.43$ | 2.3 - 6 m $p = 0.01$ |
| | | | 6 - 10 m $p = 0.001$ |
| | | | 10 - 15 m $p = 0.89$ |
| | | | 15 - 18 m $p > 0.99$ |
| | | *Post-VC*<br>αCGRP loss: $F(1, 81) = 24.82, p < 0.001$<br>Aging: $F(3, 81) = 0.4429, p = 0.72$ | 2.3 - 6 m $p = 0.004$ |
| | | | 6 - 10 m $p < 0.001$ |
| | | | 10 - 15 m $p > 0.99$ |
| | | | 15 - 18 m $p > 0.99$ |
| | Female | *Pre-VC*<br>αCGRP loss: $F(1, 79) = 1.94, p = 0.17$<br>Aging: $F(3, 79) = 2.37, p = 0.08$ | 2.3 - 6 m $p = 0.0009$ |
| | | | 6 - 10 m $p = 0.23$ |
| | | | 10 - 15 m $p = 0.59$ |
| | | | 15 - 18 m $p > 0.99$ |
| | | *Post-VC*<br>αCGRP loss: $F(1, 79) = 6.722, p = 0.01$<br>Aging: $F(3, 79) = 2.342, p = 0.08$ | 2.3 - 6 m $p < 0.001$ |
| | | | 6 - 10 m $p = 0.009$ |
| | | | 10 - 15 m $p = 0.48$ |
| | | | 15 - 18 m $p > 0.99$ |
| Postural Sway | Male | *Pre-VC*<br>αCGRP loss: $F(1, 53) = 0.882, p = 0.35$<br>Aging: $F(3, 53) = 4.673, p = 0.006$ | 2.3 - 6 m $p > 0.99$ |
| | | | 6 - 10 m $p > 0.99$ |
| | | | 10 - 15 m $p > 0.99$ |
| | | | 15 - 18 m $p = 0.32$ |
| | | *Post-VC*<br>αCGRP loss: $F(1, 53) = 0.2654, p = 0.61$<br>Aging: $F(3, 53) = 0.7518, p = 0.53$ | 2.3 - 6 m $p > 0.99$ |
| | | | 6 - 10 m $p > 0.99$ |
| | | | 10 - 15 m $p = 0.67$ |
| | | | 15 - 18 m $p > 0.99$ |
| | Female | *Pre-VC*<br>αCGRP loss: $F(1, 68) = 0.947, p = 0.33$<br>Aging: $F(3, 68) = 2.508, p = 0.07$ | 2.3 - 6 m $p > 0.99$ |
| | | | 6 - 10 m $p = 0.35$ |
| | | | 10 - 15 m $p > 0.99$ |
| | | | 15 - 18 m $p > 0.99$ |
| | | *Post-VC*<br>αCGRP loss: $F(1, 68) = 6.031, p = 0.02$<br>Aging: $F(3, 68) = 4.847, p = 0.007$ | 2.3 - 6 m $p = 0.09$ |
| | | | 6 - 10 m $p = 0.009$ |
| | | | 10 - 15 m $p > 0.99$ |
| | | | 15 - 18 m $p > 0.99$ |

**Table 3:** Analyzes were separately performed in males and females. To determine the impact of a vestibular challenge (VC) on these balance behaviors, rotarod and postural sway data were further analyzed with two-way repeated measures ANOVA to assess the factors *aging* and *VC effects* in wildtype or αCGRP (−/−) null data. Bonferroni post hoc analyses computed the differences between pre-VC and post-VC outcomes at each age group. F-values are listed in degrees of freedom (DF_n_, DF_d_) and p-values are given.

| Assay | Biological Sex | ANOVAs<br>F-statistics, p-val | Post-hoc<br>pre vs post VC \| p-val |
| --- | --- | --- | --- |
| Rotarod | Male | *Wild type*<br>Aging: $F(3, 42) = 3.756, p = 0.02$<br>VC effect: $F(1, 42) = 0.081, p = 0.78$ | 2.3 - 6 m $p > 0.99$ |
| | | | 6 - 10 m $p > 0.99$ |
| | | | 10 - 15 m $p > 0.99$ |
| | | | 15 - 18 m $p > 0.99$ |
| | | αCGRP(-/-) null<br>Aging: $F(3, 42) = 2.632, p = 0.06$<br>VC effect: $F(1, 42) = 13.50, p < 0.001$ | 2.3 - 6 m $p > 0.99$ |
| | | | 6 - 10 m $p = 0.01$ |
| | | | 10 - 15 m $p = 0.11$ |
| | | | 15 - 18 m $p > 0.99$ |
| | Female | *Wild type*<br>Aging: $F(3, 37) = 4.581, p = 0.008$<br>VC effect: $F(1, 37) = 3.418, p = 0.07$ | 2.3 - 6 m $p > 0.99$ |
| | | | 6 - 10 m $p > 0.99$ |
| | | | 10 - 15 m $p = 0.11$ |
| | | | 15 - 18 m $p = 0.62$ |
| | | αCGRP(-/-) null<br>Aging: $F(3, 42) = 4.583, p = 0.007$<br>VC effect: $F(1, 42) = 12.33, p = 0.001$ | 2.3 - 6 m $p > 0.99$ |
| | | | 6 - 10 m $p = 0.24$ |
| | | | 10 - 15 m $p < 0.05$ |
| | | | 15 - 18 m $p = 0.20$ |
| Postural Sway | Male | *Wild type*<br>Aging: $F(3, 27) = 2.86, p = 0.06$<br>VC effect: $F(1, 27) = 27.15, p < 0.001$ | 2.3 - 6 m $p > 0.99$ |
| | | | 6 - 10 m $p = 0.27$ |
| | | | 10 - 15 m $p < 0.001$ |
| | | | 15 - 18 m $p < 0.001$ |
| | | αCGRP(-/-) null<br>Aging: $F(3, 26) = 2.04, p = 0.13$<br>VC effect: $F(1, 26) = 33.41, p < 0.001$ | 2.3 - 6 m $p > 0.99$ |
| | | | 6 - 10 m $p = 0.08$ |
| | | | 10 - 15 m $p < 0.001$ |
| | | | 15 - 18 m $p = 0.006$ |
| | Female | *Wild type*<br>Aging: $F(3, 37) = 2.17, p = 0.11$<br>VC effect: $F(1, 37) = 1.20, p = 0.28$ | 2.3 - 6 m $p > 0.99$ |
| | | | 6 - 10 m $p = 0.33$ |
| | | | 10 - 15 m $p = 0.006$ |
| | | | 15 - 18 m $p = 0.07$ |
| | | αCGRP(-/-) null<br>Aging: $F(3, 31) = 2.04, p = 0.13$<br>VC effect: $F(1, 31) = 23.06, p < 0.001$ | 2.3 - 6 m $p = 0.75$ |
| | | | 6 - 10 m $p = 0.02$ |
| | | | 10 - 15 m $p = 0.01$ |
| | | | 15 - 18 m $p = 0.02$ |

### VC’s effects on rotarod

Mice were challenged by a 30-second circular rotation (see methods) and were immediately tested afterward on rotarod. Post-VC outcomes resembled pre-VC in either sex and strain (**Fig 1D**). In females and males, WT mice outperform αCGRP (−/−) null at early and late adulthood but resemble αCGRP (−/−) null performance after 10 months of age (WT vs KO (2.3 – 6 m): *p* = 0.004 for males and *p* < 0.001 for females; WT vs KO (6 – 10 m): p < 0.001 for males and p = 0.009 for females) (**Fig 1E**). To more clearly observe the effects of VC on an individual animal’s performance, before-after plots were constructed for WT male (**Fig 1F**), αCGRP (−/−) null male **(Fig 1G)**, WT female **(Fig 1H)**, and αCGRP (−/−) null female **(Fig 1I)**. Separate 2-way RM-ANOVAs were used to assess VC’s effects and aging (top half of **Table 3**), but the VC did not appear to impact performance, except for the following, particular cases: αCGRP (−/−) null males in late adulthood showed a decrease of about 4.76 + 1.50 s due to the VC (*p = 0*.*01*) and αCGRP (−/−) null females at middle age showed a decrease of 3.99 + 1.52 s (*p < 0*.*05*) due to VC. Based on this data, the effect of a vestibular challenge in bringing out aging deficits in a mouse model for dynamic imbalance is present, but is generally unclear and may require further exploration.

### Aging and αCGRP loss (−/−) on Postural Sway

In **Fig 2B**, each symbol indicates the average CoP for a particular animal. As similarly done for rotarod, 2-way RM-ANOVAs on CoP were computed across the factors αCGRP loss and aging during pre-VC and post-VC tests, and these analyses were done separately in males and females (bottom half of **Table 2)**. Group average CoPs + SEM are depicted in **Fig 2D**. During the pre-VC test, we observed a general increase in CoP area from early adulthood to middle age, but no significant differences were observed between WT and αCGRP (−/−) null during the pre-VC tests.

### VC’s effects on Postural Sway

Mice were assessed for postural sway as measured by center of pressure (CoP) changes after a 5-minute orbital rotation (see methods). In females only, WT exhibited larger CoP ellipses at early adulthood when than their αCGRP (−/−) null complement (t (*p* = 0.09)). This was also observed at late adulthood between WT and αCGRP (−/−) null females (*p* = 0.0009). However, this increased CoP observation is gone as the WT females age past 10 months. From this point onward, no differences were observed between strains at middle age and senescence. Separate 2-way RM-ANOVAs assessed VC’s effects and aging on CoP outcomes and are depicted in the bottom half of **Table 3**. For WT female (**Fig 2F**), post-VC CoP ellipse areas were decreased at middle age by 1.32 + 0.38 cm^2^ (*p = 0*.*006*) and at senescence by 1.27 ± 0.51 cm^2^ (t (*p = 0*.*07*)). A similar result was observed in WT males (**Fig 2G**), as post-VC ellipse areas were decreased at middle age by 1.34 + 0.31 cm^2^ (*p < 0*.*001*) and at senescence by 1.27 ± 0.29 cm^2^ (*p < 0*.*001*). Interestingly, the αCGRP (−/−) nulls showed a response to VC earlier than WT. For αCGRP (−/−) null females (**Fig 2H**), post-VC CoP ellipse areas were decreased in late adulthood by 1.12 ± 0.38 cm^2^ (*p = 0*.*02*), at middle age by 1.27 ± 0.40 cm^2^ (*p = 0*.*01*), and by 0.90 ± 0.30 cm^2^ (*p = 0*.*02*). For αCGRP (−/−) null males (**Fig 2I**), post-VC CoP ellipse areas were decreased in late adulthood by 0.65 ± 0.27 cm^2^ (*t (p = 0*.*08)*), at middle age by 1.45 ± 0.266 cm^2^ (*p < 0*.*001*), and by 0.85 ± 0.24 cm^2^ (*p = 0*.*006*). In addition, mice with significantly low CoP ellipse areas during the post-VC test exhibited a “freezing behavior”, characterized by a lack of activity and a fixed gaze. This behavior is distinct from the usual behavior observed by these mice; however, it was not further quantified in this study.

## Discussion

The vestibular system’s ability to maintain balance and coordination deteriorates as a function of aging. Previous findings show aging’s detrimental effects on balance control in humans and rodents can arise from changes in the vestibular endorgans and vestibular nerve [16–19], chemical imbalances in the inner ear [20], and age-related musculoskeletal deterioration [21, 22]. Mouse models provide an opportunity to assess aging’s effects on behavior when an essential neuromodulator like CGRP is removed from birth. The αCGRP (−/−) null mice are characterized with a reduced efficacy of the vestibulo-ocular reflex [11], an enhanced activation timing of primary vestibular afferents, a lack αCGRP expression in otolith organs, and a poor rotarod ability at an early age [12]. In addition to CGRP’s role as an efferent neurotransmitter; CGRP is present in the vestibular nuclei and the cerebellum [23, 24], and losing CGRP from birth is hypothesized to strongly affect the upstream pathways to these structures that ultimately integrate vestibular and motor signals. Thus, we were interested in assessing aging’s effects on the balance behavior of αCGRP (−/−) null mice..

Rotarod performance is a mouse surrogate for dynamic imbalance. Our results show that WT mice perform better on the rotarod than αCGRP (−/−) null mice from 2.3 to 10 months, but the difference disappears after 10 months. Our results in the WT correlate with a previous report assessing rotarod and balance beam on the aging C57BL/6 mouse tested at one, nine, and thirteen months of age [25]. In this prior study, a reduction in rotarod performance was observed at nine months and longer latencies to traverse a balance beam were measured at nine and thirteen months of age. Comparatively, the αCGRP (−/−) null mice have poor gait and dynamic balance early in life but do not experience further decline or noticeable improvement. Interestingly, CGRP(−/−) mice are hypothesized to have impaired spatial learning due to lower Insulin-Like Growth Factor-1 levels compared to WT, and thus a cognitive impairment may be one justification for an inability to do well on rotarod [26]. We did not observe a significant impact of the vestibular challenge (VC) on rotarod performance for either strain, as this lack of VC effect on rotarod was also observed in the aging C57BL/6 study. This other study’s VC led to increased balance beam latencies in the C57BL/6 mice, and would be an interesting future direction for research in the αCGRP (−/−) null. It is known that the organism’s vestibular system adapts quickly to changes in its environment, and mice can recover from the vestibular challenge almost immediately [25]. This quick recovery provides a rationale as to why we detected no significant impact of the vestibular challenge on max LTFs (a given mouse’s best effort on the rotarod), and highlights the adaptability of the vestibulo-muscular systems in responding to brief but intense vestibular perturbations.

In postural sway testing, we observed WT females exhibited larger CoP sway ellipses in both early and late adulthood than compared to their αCGRP (−/−) null complement and to WT and αCGRP (−/−) null males. However, this increased CoP observation in WT females is no longer present as mice age past 10 months. In response to the vestibular challenge (VC), mice start to experience a “freezing” behavior characterized by a lack of movement and a fixed gaze. This behavior is associated with significantly reduced CoP ellipse areas and typically occurs when mice are 10 months and older. Prior studies in humans have reported this freezing behavior when assessing postural control and heart rate changes to threatening stimuli [27, 28]. A stiffening strategy is observed in response to vestibular threats to one’s standing posture, and involves active engagement of the lower body muscles [29].

## Conclusions

We conclude that aging, vestibular challenge, and the loss of CGRP all contribute to deficits in both static and dynamic balance control. These imbalance deficits begin to appear at ∼10 months in the aging mouse.

## Acknowledgements

This work was supported by NIH R01-DC017261.

